# A novel 5D brain parcellation approach based on spatio-temporal encoding of resting fMRI data from deep residual learning

**DOI:** 10.1101/2021.04.22.440936

**Authors:** Behnam Kazemivash, Vince D. Calhoun

## Abstract

**Objective:** Brain parcellation is an essential aspect of computational neuroimaging research and deals with segmenting the brain into (possibly overlapping) sub-regions employed to study brain anatomy or function. In the context of functional parcellation, brain organization which is often measured via temporal metrics such as coherence, is highly dynamic. This dynamic aspect is ignored in most research, which typically applies anatomically based, fixed regions for each individual, and can produce misleading results.

**Methods:** In this work, we propose a novel spatio-temporal-network (5D) brain parcellation scheme utilizing a deep residual network to predict the probability of each voxel belonging to a brain network at each point in time.

**Results:** We trained 53 4D brain networks and evaluate the ability of these networks to capture spatial and temporal dynamics as well as to show sensitivity to individual or group-level variation (in our case with age).

**Conclusion:** The proposed system generates informative spatio-temporal networks that vary not only across individuals but also over time and space.

**Significance:** The dynamic 5D nature of the developed approach provides a powerful framework that expands on existing work and has potential to identify novel and typically ignored findings when studying the healthy and disordered brain.

## I. Introduction

**T**HE brain is incredibly complex and the most sophisticated organ in the body, comprised of copious neurons collaborating precisely to control perception, emotion, intelligence, creativity and general day-to-day human life activities [6]. It is composed of different anatomic subparts including cerebrum, cerebellum and brainstem which are anatomically linked and functionally coupled to conduct a specific cognitive function or produce a behavioral response to external stimulus [5,7].

Computational neuroimaging is an interdisciplinary research domain that deploys many fields including visual neuroscience, computer science and psychology for investigating brain organization and function and for developing computational theories [6,9]. Brain parcellation as a branch of computational neuroimaging seeks to segment the brain into a mosaic of anatomic or functionally distinct regions which are spatially homogeneous [36, 40]. Brain parcellation has a significant role in exploring the connectome, a comprehensive brain map of neural connections, and can be utilized as a prior for studies of brain disorders [8,6].

Due to the natural diversity of brain shape and cortical folding (gyrification), as well as the cytoarchitectonic variations which are not visible via macroscopic imaging approaches like magnetic resonance imaging (MRI), proposing an efficient and rigorous parcellation scheme which reflects the functional and structural variability among individuals can significantly advance the field [10, 37]. Existing parcellation approaches can be divided into atlas (anatomy)-based and connectivity (function)-based approaches [11]. Atlas-based approaches are widely used and typically segment the brain by applying a predefined anatomical template on the input image and generating a set of regions over associated template. Despite the popularity of these methods, there are drawbacks [14, 39]. Obviously, these methods rely heavily on the given template and do not take natural variance of the brain into account for different subjects beyond the geometric warping of the fixed atlas into a standardized space using either voxel-based or surface-based approaches. The spatial registration procedures exploited in these techniques can be computationally expensive [12, 15].

Connectivity-based methods, on the other hand, segment the brain into temporally coherent portions of homogeneous patterns considering similarity metrics. Choosing a suitable similarity metric is a critical step in connectivity-based methods and can significantly affect the result [13, 18]. Independent component analysis (ICA) is a type of connectivity-based approach [3]. ICA is widely used and provide a source-based technique that can adapt to the data at hand. ICA-based methods adapt to an individual brain, but typically assume a fixed spatial pattern for each network over time within a given person. Changes over time are captured by a single timecourse which modulates the contribution of the network to each point in time. Furthermore, ICA-based approaches commonly exploit free parameters such as the number of components which can be used to model the brain at different spatial scales [4]. Another advantage of ICA, relative to most other parcellation approaches, is the components have values at all voxels, thus allowing for overlap. This facilitates modeling of voxels which participate in multiple functional networks.

There is a considerable amount of research studying functional connectivity and brain dynamics for estimating and visualizing synchronized interactions of focal or distributed brain regions through time and linking of these to cognition and behavior in the healthy and disordered brain [16, 41]. There are also many different ways to model brain dynamism including spatial, temporal, and spatio-temporal dynamics, each of which provide unique and relevant information on brain activity and connectivity [17]. Temporal dynamic approaches to brain connectivity deals with studying time-varying properties of functional magnetic resonance imaging (fMRI) data by evaluating transient changes in temporally correlated or mutually informed information between fixed brain regions or spatial nodes [19]. More recently there has been recognition the need for also modeling changes in the spatial nodes/networks over time as well [17, 35] including changes in shape, size or translation of active regions. If we consider both spatial and temporal features for a given subject simultaneously then spatio-temporal dynamics is taken into account [38].

While abundant research has been conducted on brain parcellation, to our knowledge, none of them has modeled the full 4D nature of human brain function, allowing for both temporal and spatial changes over time. The computational complexity for such an approach can be challenging, but the improved efficiency of machine learning algorithms provides a substantial opportunity. We herein propose a novel multi-model system which adopts deep residual network structure to predict brain networks for fMRI images which can be used for tracking spatio-temporal dynamics over space, time, and network, resulting in a 5-dimensional parcellation (i.e., 4D spatiotemporal information for each brain network).

The remaining text is organized as follows. Section II reviews related scientific literature followed by theoretical background in section III. Also, section IV discusses details of the proposed method and continues with results in section V. Next, we have discussion section VI to analyze experimental results and finally, we have conclusion in section VII.

## II. related work

The idea of employing predefined templates in brain parcellation or taking advantages of mathematical similarity metrics has been widely investigated in the neuroimaging and neuroscience community. Machine learning algorithms being applied in brain parcellation have shown good results [20]. Recently, Liu et al. presented a new multi-atlas adversarial fully convolutional network (FCN) [21] for brain parcellation in which the generative model is an FCN merged with brain atlases information and the discriminative model is a convolutional neural network with multi-scale deep features. Bonmati et al. implemented a novel and efficient framework based on information theory that models brain regions as a random walk on the connectome by applying a Markov process in which the different nodes refer to brain regions. They also used an agglomerative information bottleneck technique to cluster functional or anatomical patches by minimizing loss of information through computing mutual information as the similarity metric [24].

In another approach, Thyreau et al. proposed a novel and efficient scheme bridging atlas-warping techniques and the precision of landmark localization in a deep learning model, leading to meaningful and interesting results [22]. Moreover, Luo et al. conducted research adopting an eigen clustering technique for brain parcellation utilizing the precuneus–cortical functional connectivity derived from fMRI images and showed that it is robust to noise [25].

In other work, Zhao et al. proposed a two-step parcellation scheme which maps resting fMRI (rs-fMRI) signals to cortical surface and then registers that into standard space. For the second step, the system estimates the similarity between vertices based on the correlation of rs-fMRI time series. This approach has shown acceptable results based on standard metrics [23]. Similarly, Boukhdhir et al. presented a novel scheme to detect dynamic states of segments; In their method, a clustering analysis has been executed to regroup short time windows into states with similar seed patches [26].

In addition to studying relevant work, Graham et al. [27] proposed a novel brain parcellation scheme by generating a hierarchical classification tree which is adapted from a segmentation task and extending it for predicting decisions (by) modeling an uncertainty factor for each of them. However, none of the above approaches model the full 4D aspect of brain functional connectivity.

## III. theoretical Background

Our brain parcellation scheme exploits ICA output to extract components that are used as prior information to train a deep residual model in a supervised manner. ICA extracts maximally spatially independent maps from the fMRI data and their associated time courses. The data, for a given fMRI dataset *X* which is time-by-space, are decomposed into a time course matrix *A* which is time-by-component and the spatial maps *S* which is component-by-voxel, written as

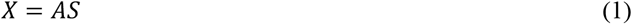

This can also be done at the group level as in [2]. We leverage a recently proposed fully automated spatially constrained ICA pipeline called NeuroMark [1] which enables us to estimate brain functional network from functional magnetic resonance imaging (fMRI) data that can be used to link brain network abnormalities among different datasets, studies, and disorders. As mentioned, the ICA decomposition does not explicitly model temporal variation of the spatial maps. We can utilize this pipeline however by taking the spatial maps as priors to initialize the model.

## IV. method

We propose a novel predictive model that encodes spatio-temporal dynamics to estimate individual subject brain network parcellations that are linked across subjects as well. Accordingly, we frame the approach as a regression problem by predicting the probability of each voxel belonging to different brain networks. Each brain network is derived from a combination of temporal and spatial ICA components computed for a given subject using the approach discussed in the theoretical section. Next, we train a deep residual model for each of the brain networks via a supervised approach by using the fMRI time series images as input and generating labels as the target. In terms of novelty, the proposed scheme utilizes a deep learning structure to encode a set of 4D fMRI images into network specific brain parcels which vary through time as shown in figure 1. To do this, for each network we apply the pre-trained model to each timepoint in a repetitive manner to achieve the entire 4D network. We do this for all networks, resulting in a full 5D parcellation of the brain.

**Fig. 1.**
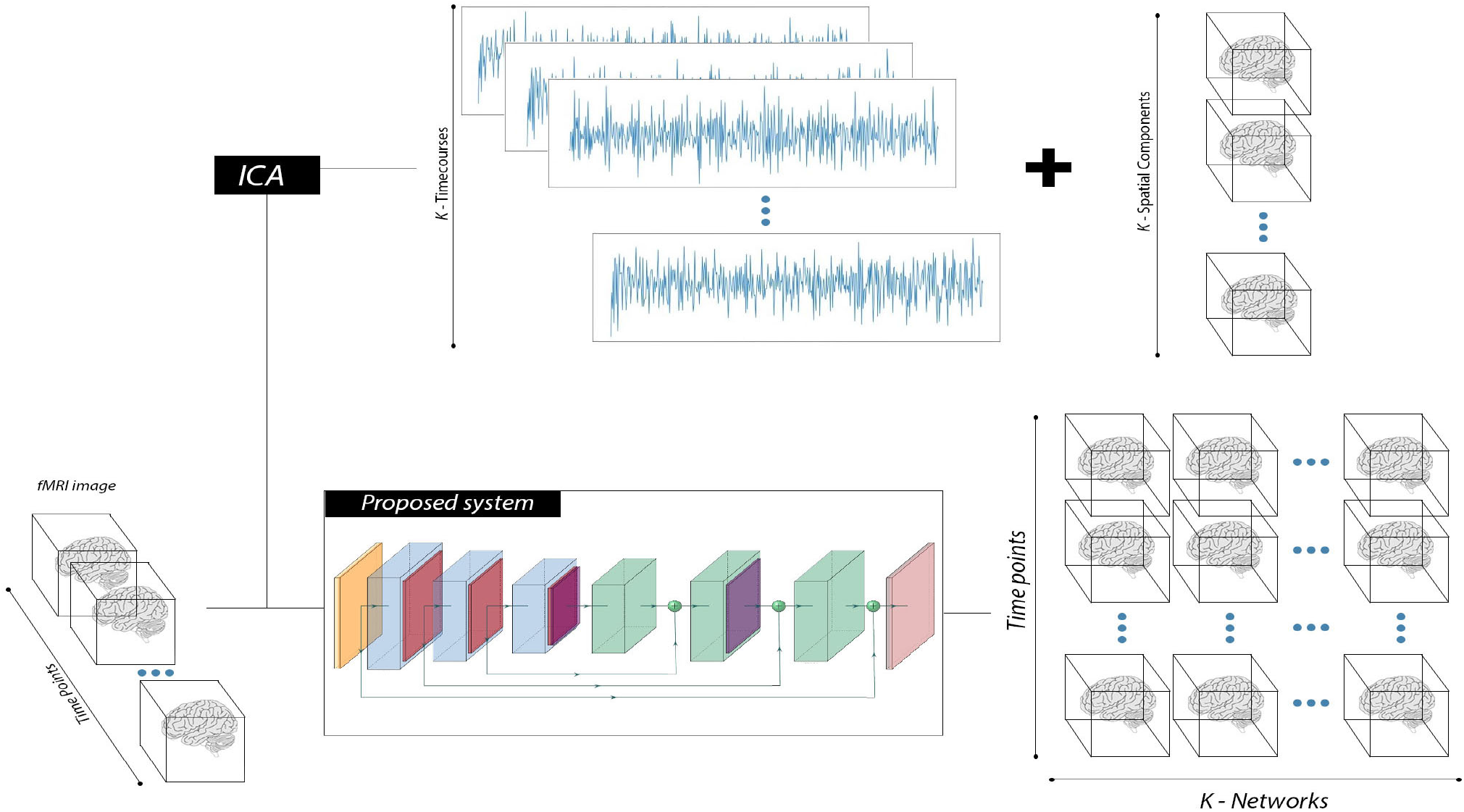
General concept of ICA-based methods versus the proposed approach.

### A. Label Generation

Generating labels is a critical step in proposed approach due to our supervised model training. First, we applied ICA on fMRI images each of which are a 4D tensor with one temporal axis and 3 dimensions for indicating spatial coordinates. By applying ICA and extracting components, we have *k* time courses and spatial maps separately where a time course is a vector with size *t* equal to the length of time axis in the original fMRI image, and a spatial map is a vector of which the size is product of spatial coordinates, the voxel size. Then, we compute the outer product of time courses and spatial maps for all *k* components.

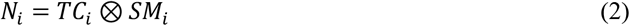

where, *TC_i_ is* a specific time course, *SM_i_* refers to a given spatial map and ⊗ is the outer product operator. Clearly, *N_i_* is a matrix with size of *t*-by-*voxel size*. We then build up a 3D tensor by repeating the procedure for all *k* components. Next, we generate probabilistic maps by applying the following equation:

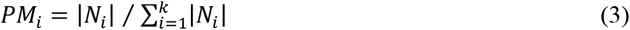

Here, *PM* shows the probabilistic map, and the dominator is the elementwise summation of absolute values of *N* which forms a matrix with same shape as *N.* This procedure guarantees all elements range between 0 and 1 because we divide each element with norm-1 of a vector that includes different values of each elements in *k* maps. Finally, we transform each probability network into a 4D tensor by deploying a predefined mask with the same shape as the input fMRI image.

### B. Model Configuration

Our regression model is configured as a deep residual network with 36 layers including conv3D, transposed Conv3D, batch normalization, max pooling and drop out layers which are grouped as encoding and decoding blocks and finally skip connections inside and outside of each block. The model structure has 2 focal sections of encoder and decoder sections; The encoder begins by a conv3D layer with output channel size of 64 and kernel size of 3 followed by sigmoid function. Next, we have residual encoding block with 3 volumetric convolution and batch normalization layers with same out-channel size of 32 and also both of last conv layers use padding size of 1. Also, we apply sigmoid function on each couple of conv and batch norm layers’ output.

Furthermore, we have a 3D max pooling layer just after each encoding block with stride of 1 and same kernel size as convolution layers. The remaining encoding blocks have identical configuration and dependencies as first block except out-channel size of 16 and 8. The last layer in encoder section of model is a 3D dropout layer with ratio of 0.5. On the decoder side, we have 2 consecutive decoding block each of which has a couple of 3D conv transposed layers followed by batch normalization layers and a sigmoid activation function immediately after this layer. The sizes of the out-channels are, respectively, 16 and 32.

We also have a dropout layer with ratio of 0.5. Subsequently, there is another decoding block with an out-channel size of 64 and similar remaining parameters. The last layer is a 3D conv transposed layer with the out-channel size of 1 and kernel size of 3. Furthermore, we applied mean squared error as a loss function in the supervised training procedure as shown in the equation below:

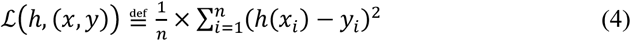

The schematic diagram of the model structure contains encoding and decoding blocks along with skip connections, as shown in figure 2.

**Fig. 2.**
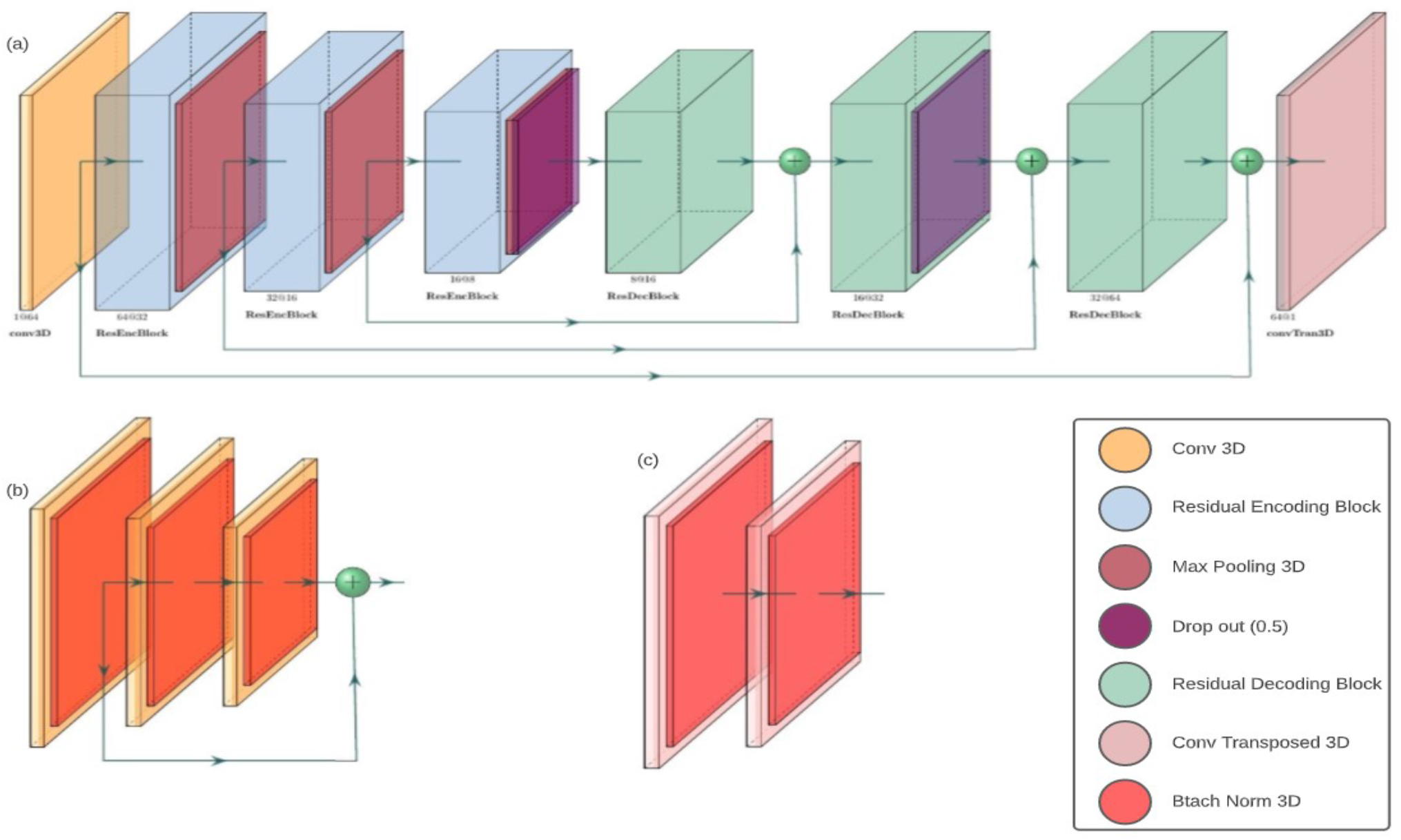
Schematic diagram of model organization with detail structure of (b) encoding block and (c) decoding block.

### C. Model input templates

For the spatial priors, we utilize the NeuroMark templates [33]. A template of replicable independent components (ICs) constructed after spatially matching correlated group-level ICs between two large (N>800) healthy control fMRI datasets, genomics superstructure project (GSP) and human connectome project (HCP). The template was then used as priors for a spatially constrained ICA algorithm applied to each subject individually to identify 53 functionally relevant resting-state networks (RSNs) for each individual that are maximally spatially independent. Each RSN consists of one spatial map, with a size of 53×63×52 voxels, and its associated time course. RSNs are grouped into seven domains, namely subcortical (SC), auditory (AU), sensory motor (SM), visual (VI), cognitive control (CC), default mode (DM), and cerebellar (CB), by function similarity.

### D. fMRI data: Acquisition and preprocessing

To evaluate preliminary results, we applied our approach to data from the UK Biobank study (six subjects to evaluate the ability of our approach to capture variability within subjects, and forty subjects to evaluate the ability of our approach to capture changes related to a subject variable, age). Participants were scanned once by a 3-Tesla (3T) Siemens Skyra scanner with a 32-channel receive head coil, acquired all in one site. A gradient-echo echo planar imaging (GE-EPI) paradigm was used to collect/obtain resting-state fMRI scans. The EPI-based acquisition parameters include multiband acceleration factor of 8 (i.e., eight slices were acquired simultaneously), no iPAT, fat saturation, flip angle (FA)=52°, spatial resolution = 2.4×2.4×2.4mm, repeat time (TR)=0.735s, echo time (TE)=39ms, and 490 volumes. Subjects were instructed to stare at a crosshair passively and remain relaxed, not thinking about anything, during the six minutes and ten second resting-state scanning period.

The preprocessing steps performed by UK Biobank are as follows. An intra-modal motion correction tool, MCFLIRT [31], was applied to minimize the distortions due to head motions. Grand-mean intensity normalization scaled the entire 4D dataset by a single multiplicative factor to facilitate comparability of brain scans between subjects. The data were filtered by a high-pass temporal filter (Gaussian-weighted least squares straight line fitting, with σ = 50.0 s) to remove residual temporal drift. Geometric distortions of EPI scans were corrected by using FSL’s Topup tool [32]. EPI unwarping is followed by a gradient distortion correction unwarping phase. More details on the imaging protocol and preprocessing steps can be found in [30].

## V. results

We implemented the entire system including the model, preprocessing and post-processing procedures in python 3.6 mainly using the Pytorch [28], NumPy and Nilearn libraries. All experiments executed on a TESLA V100 GPU with 32 GB of dedicated RAM. We trained 53 models each of which grouped into a specific domain including cerebellar, sensory-motor, default mode, auditory, subcortical, cognitive control and visual networks. Our model benefits from the Adam optimizer with an adaptive learning rate of 0.00001 and step size of 5 for optimizing the loss function and 200 epochs for training phase. We also used an early stopping technique to avoid overfitting and to improve the generalization of the trained model.

Furthermore, to evaluate the ability of our approach to capture within and between subject variability, we trained our model using data from 3 fMRI subjects from the UK Biobank’s dataset [29] each of which has 490 timepoints; A total of 1470 volumes with dimensions of (53,63,52) were normalized using a min-max method. We split the dataset into training, validation and test dataset by grouping 70, 10 and 20 percent of the entire dataset, respectively.

Finally, we trained each of 53 models by feeding input fMRI images and relevant probability network, extracted from ICA as priors based on described configuration. Moreover, training and validation losses converged to low values after primary fluctuations as shown in Figure 3 for a sample subcortical network. Consequently, the trained models show consistent behavior facing unseen samples due to the average mean squared error for test set which ranges between 29 to 33 for all networks and are located in reasonable range, although some of them are a bit higher than other ones as a consequence of applying same model parameters for all of them. We can easily obtain a similar error range for all networks by applying specific learning and decay rates for each of them separately.

**Fig. 3.**
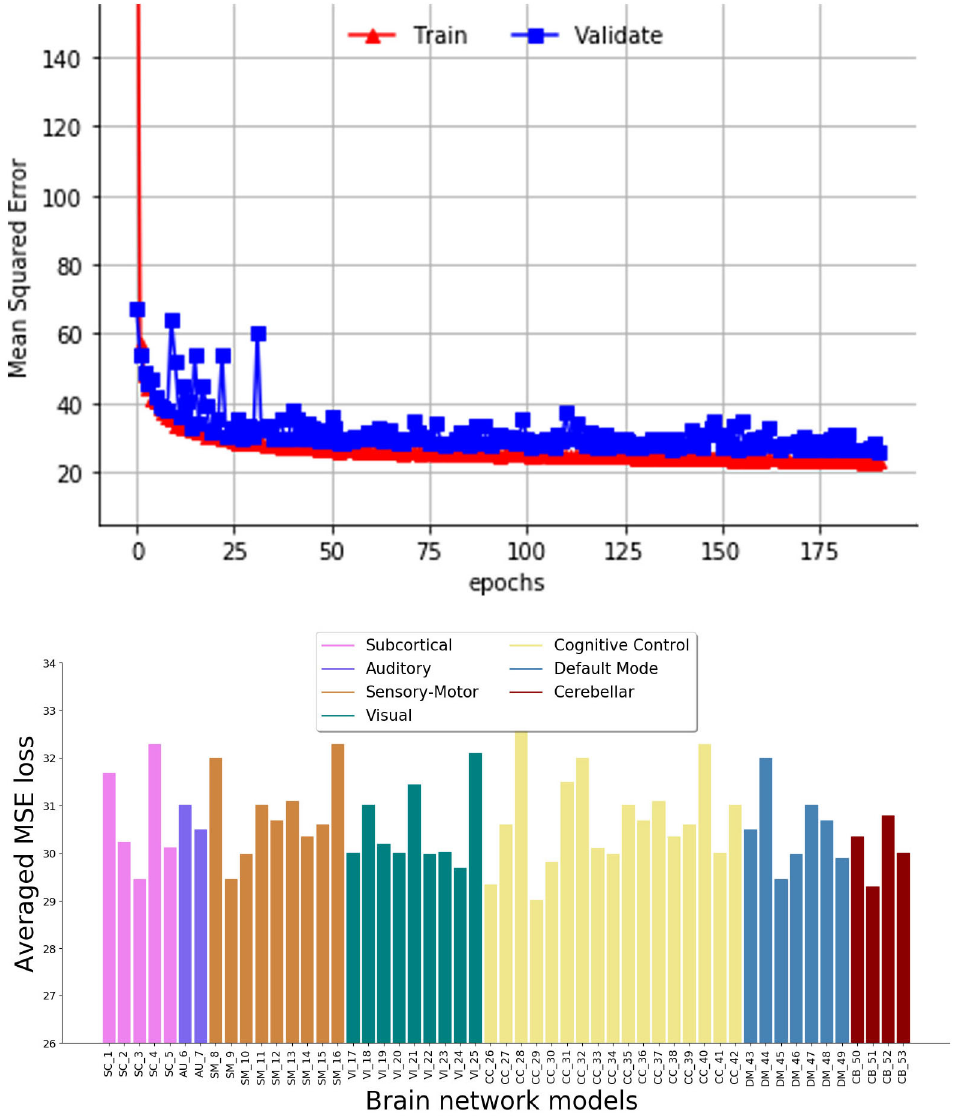
The upper plot shows the Mean Squared Error loss on training and validation set for a sample subcortical network which emphasizes on the convergences of model loss for both errors. Moreover, both errors are minimized by desirable low values. Furthermore, lower plot depicts the averaged MSE loss on test set for all 53 models each of which is related to a specific brain network. However, some of models have higher averaged MSE than others that’s mainly consequence of applying same learning rate and also decay rate for all of them instead of specific parameters but all of models’ averaged losses fluctuate in a reasonable range.

Results suggest the proposed multi-model system shows meaningful parcels, not only facilitating comparison among different subjects but also highlighting individual variability in decompositions. For example, comparing the output for one of the trained models for different subjects, we show the sensory-motor network in figure 4, which clearly emphasizes the fact that the model captures variability among subjects in the networks (in contrast to fixed atlas-based approaches); We can see smooth changes around active brain area on the output of a sensory-motor network for different subjects at timepoint 100 in figure 4 and it is worth mentioning that we also see variation along time as well, that is the spatial patterns vary within an individual over time. Moreover, the magnitude of active region varies from subject to subject or even from time point to time point that indicates ability of the trained model to capture dynamics.

**Fig. 4.**
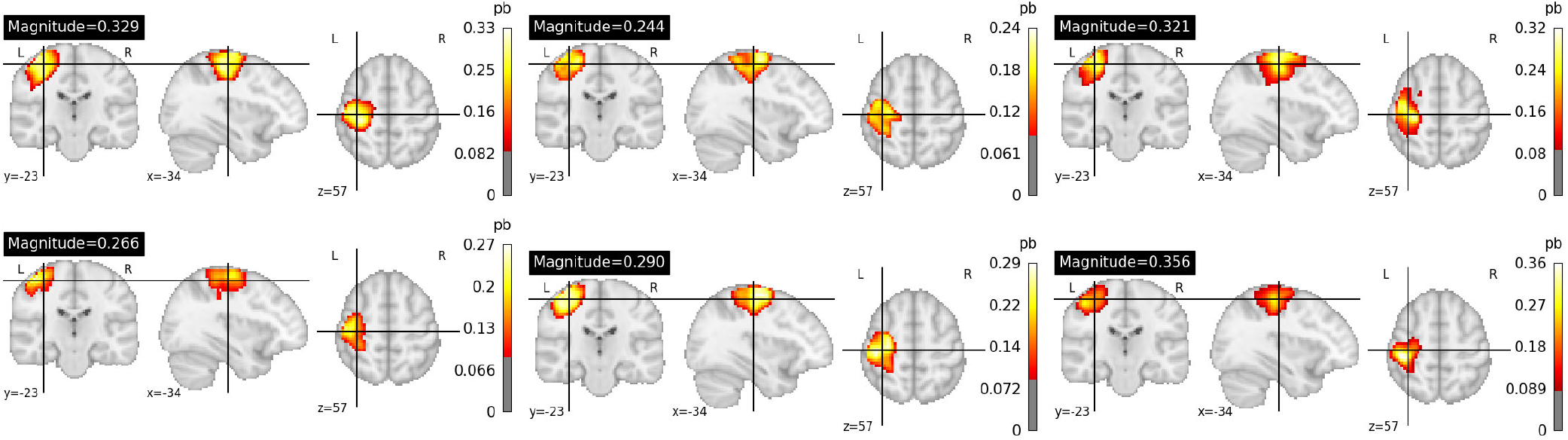
Initial differences on the output (3D image is the mean over time within a 4D sensory-motor network) for 6 different subjects on a random timepoint, highlighting the ability of our approach to capture individual variability. The “pb” unit in the figure denotes probability of each voxel belonging to this brain network.

Generally, our trained models show remarkable performance and reasonable results which can be utilized for studying healthy or disordered brain function. We depict the output of all 53 networks for a random subject by computing the average of timepoints and applying z-score to have better representation in figure 5. As is clear from the output, all trained models have extracted biologically reasonable brain regions for each of the brain networks as expected.

**Fig. 5.**
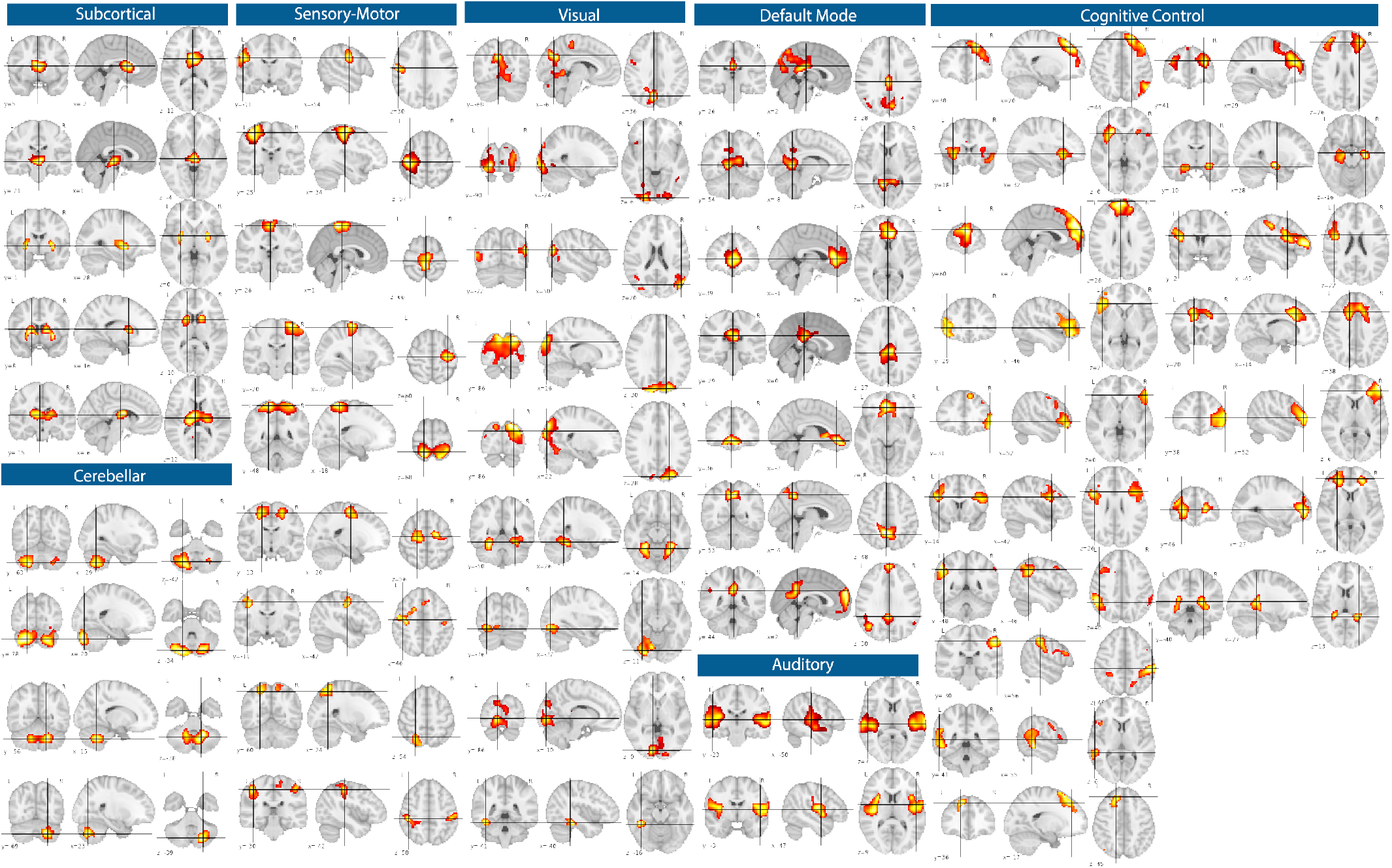
The model output for all 53 brain networks including subcortical, auditory, sensory-motor, visual, cognitive control, default mode and cerebellar networks given a sample subject. differences on the output of a sensory-motor network for 6 different subjects.

### A. Spatial Variability

One of the important features and goals of the proposed approach is to capture spatial variability over time within each network for different subjects. We next summarize the spatial variability feature in three ways. First, we processed fMRI data with our model and applied z-scoring on the model output. Next, we computed timepoint-wise differences of the previous output and summed up these differences, to capture variability over time within each voxel of a given network. Finally, we apply this to the multiple networks. We also compute the absolute timepoint-wise differences to evaluate the regions which are showing higher or lower variability regardless of sign.

The spatial variability for the sensory motor, default mode, auditory, subcortical, cerebellar and visual networks is show in figure 6. One important observation is we see that spatial variability is highlighting regions that fall within or near the thresholded network boundaries on the left but are different from those of the overall mean activity, showcasing a key advantage of our approach to provide additional insights into brain activity. For example, the absolute variation in the sensorimotor network highlights the sensorimotor strip, most of which is not captured by the amplitude (presumably because it fluctuations between positive and negative connectivity which averages to a low value over time). This highlights the additional information that can be captured when accounting for spatial variation. Another interesting example, the default mode network show very high variability in the posterior cingulate, but not in the anterior (medial frontal) portion of the network. Regarding variation, the auditory network shows two distinct clusters showing positive and negative variation. These are only a few observations which highlight the (usually ignored) information that can be captured by our proposed approach.

**Fig. 6.**
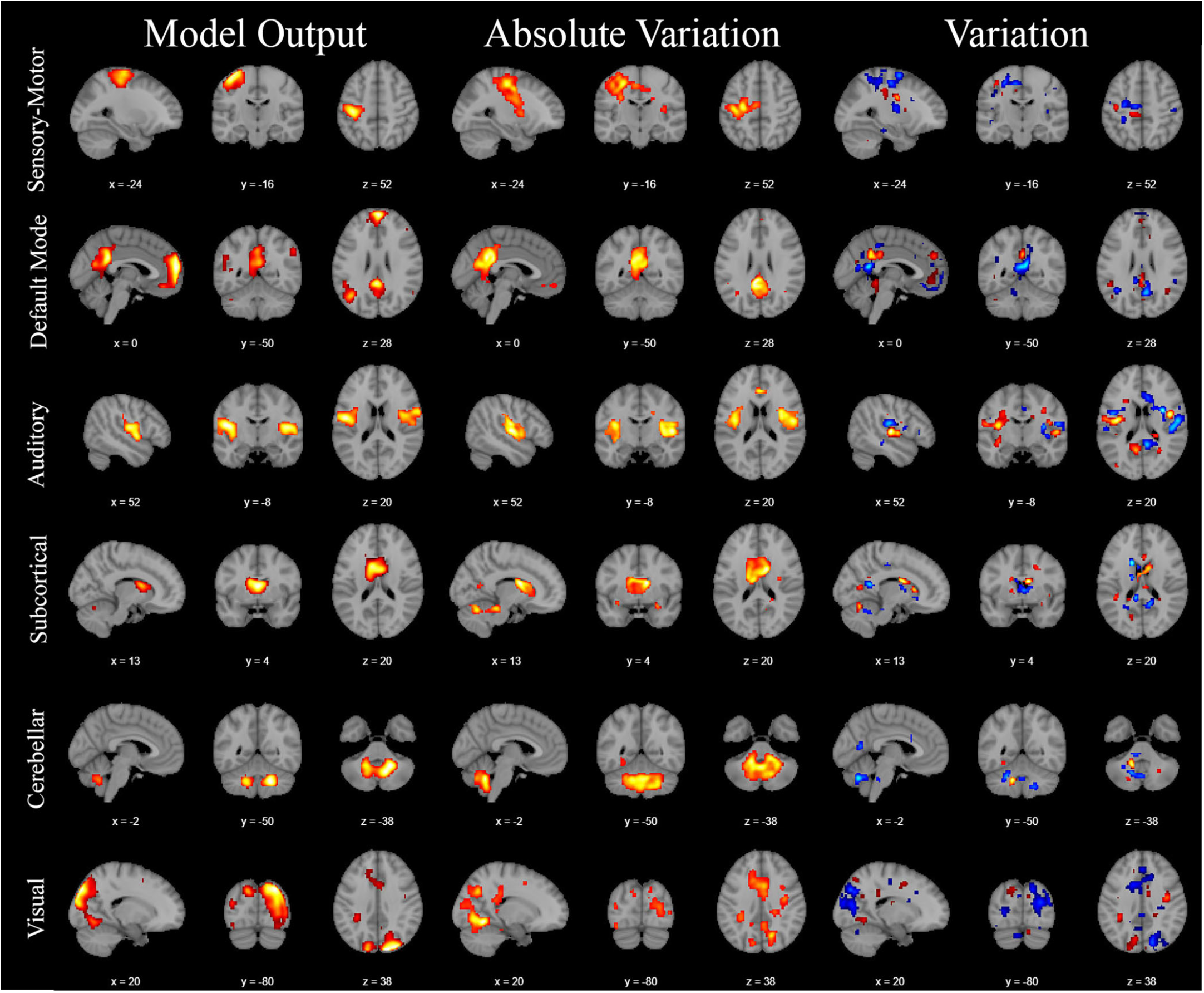
Spatial variability in sample subcortical, visual, auditory, cerebellar, default mode and sensory motor networks for a given subject. First column shows the model output (3D mean amplitude), second column is the sum of the absolute differences in time and third one shows the sum of the differences in time. All variation plots are in the same coordinates with original model output to make a fair comparison. Interestingly, the spatial variability is in some cases, most informative in regions that are highly relevant to the given network domain, but which are not captured in the average output.

### B. Age Effects

Furthermore, to study the ability of the model to capture group differences, we explored the capability of our trained models to capture age differences between subjects. To do this, we selected 20 young subjects who were less 40 years old and 20 old subjects older than 60 and computed output of the model for all of them. To summarize the captured spatial dynamics within one network (the sensorimotor network), we applied a k-means algorithm with 5 segments as the primary number of clusters using all 40 outputs. This provides us with groups of voxels which show similar spatial variability. Next, we measured the magnitude of each cluster for each subject and computed the group differences between the old and young groups for each cluster. Results showed that young subjects have higher magnitude than older subjects in all clusters.

The clustered regions for the sensory-motor network along with magnitude of the cluster showing the largest age difference (pink color) is shown in figure 7. Note that clusters with pink, green, yellow, black and blue colors, respectively, include voxels with higher to lower probability values with voxel sizes of 857, 203, 408, 402, 34860.

**Fig. 7.**
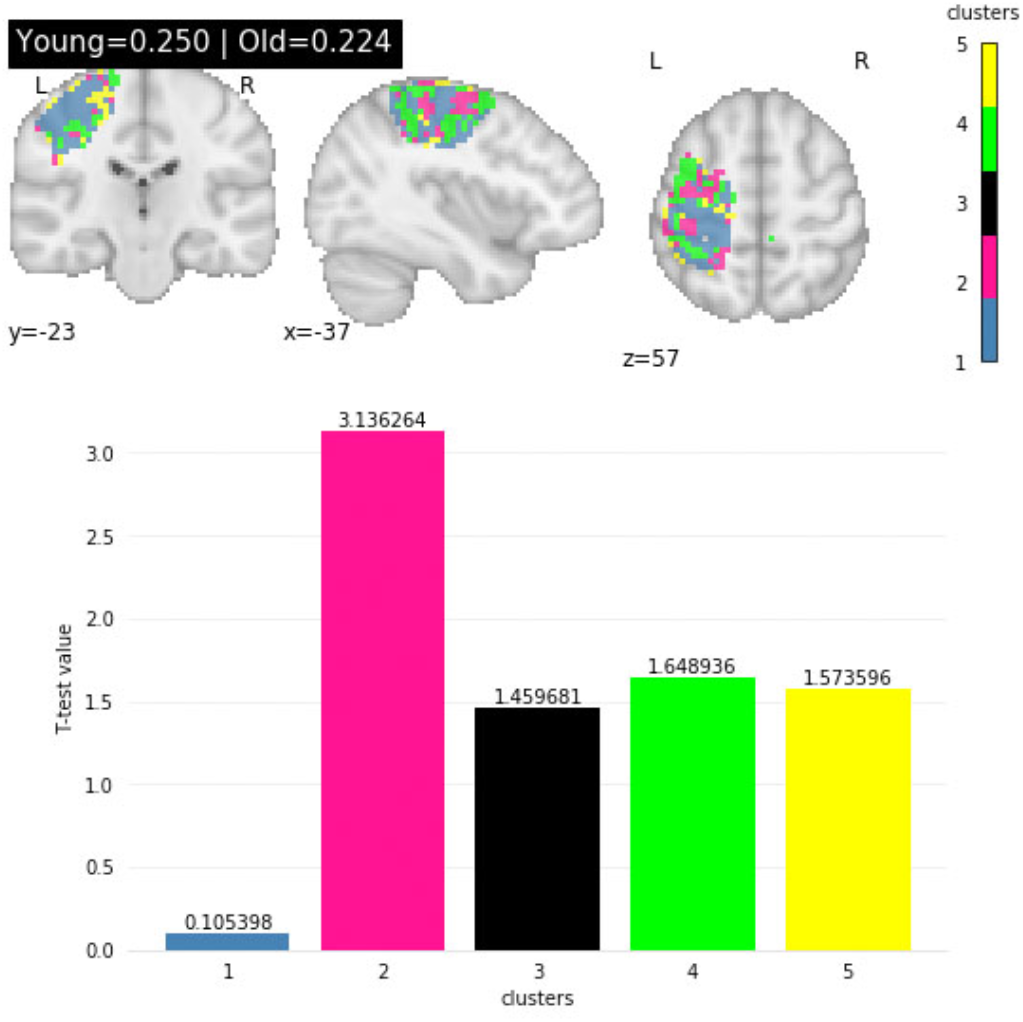
The upper figure shows clustered regions for output of a sensory-motor network after applying a threshold to study age effects on young vs old groups of subjects. All clusters are marked using different colors and pink, green, yellow, black and blue clusters, respectively, contain voxels with higher to lower probabilities. The lower figure indicates the two-sample T-Test score for young vs old subjects for all clusters. Results show that the T-value for young vs old group difference shows higher mean values for younger subjects with significant differences in the pink cluster (p<0.004).

Furthermore, we computed the two sample T-Test score for each cluster as a metric to evaluate differences in model output for young versus old subjects as is shown in figure 7. The pink cluster showed a significant difference between young and old (p < 0.004) whereas the lowest score belongs to the blue region, comprising the main and largest portion of the network. Important the magnitude of these change is quite variable as are the spatial locations of the changes, which tend to focus on the boundaries of the network. Such information would be completely ignored in a typical approach that does not focus on spatial dynamic information. Clearly, this experiment highlights ability of the model to capture changes in spatial variability in different types of groups like young and old subjects.

### C. Temporal Variability

The proposed approach also enables us to study functional dynamicity in the activity of brain networks by encoding temporal variation through different time-points. This is a salient property that can open up wide range of new studies to characterize the full spatiotemporal information in these brain networks and use them to study the healthy and disordered brain. Hereupon, we have studied preliminary evidence of temporal variation on the model’s output by computing magnitude, mean and standard deviation of active regions and show they exhibit significant fluctuations over time as is shown in figure 8 for a sample visual network.

**Fig. 8.**
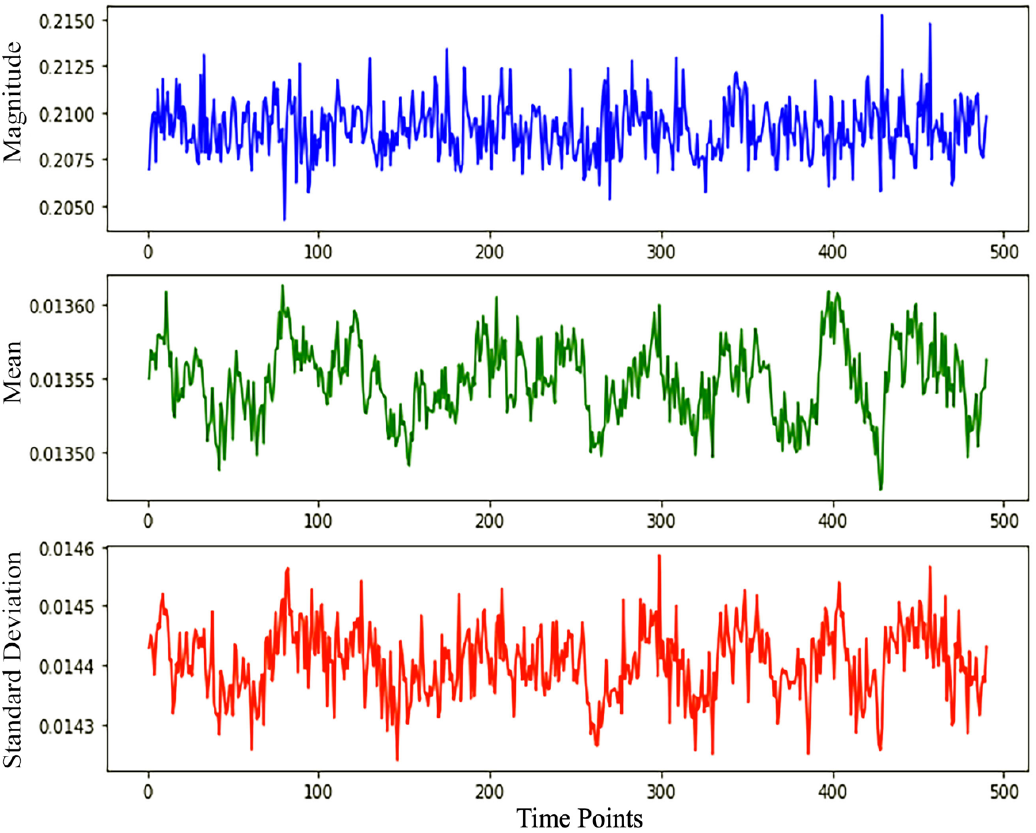
The figure shows the oscillations of magnitude (blue), mean (green) and standard deviation (red) of active region in a visual network for a given subject over 490 time points

Another approach typically used in ICA methods is to measure temporal variability or changes over time in the temporal coherence among voxels, which can be captured with metrics such as functional network connectivity (FNC) [42]. To do this, we grouped all networks into subcortical, auditory, sensory-motor, visual, cognitive control, default mode and cerebellar categories and then computed the cross correlation of all 53 networks over all timepoints and show the result as a heatmap in figure 9.

**Fig. 9.**
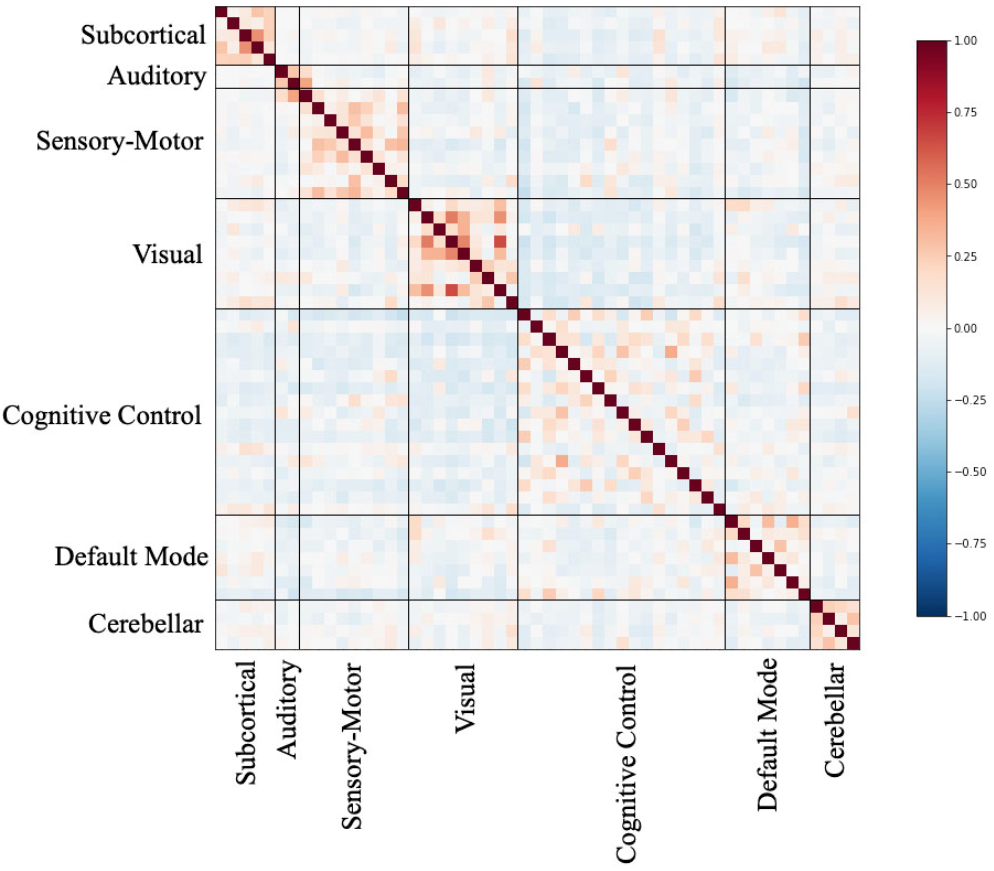
The heatmap shows the correlation between all 53 networks grouped into Subcortical, Auditory, Sensory-Motor, Visual, Cognitive Control, Default Mode and Cerebellar categories for a given subject (Static FNC).

The results exhibit the expected modularity, with generally higher correlation between networks in the same group like network 44 and 46 belonging to the default mode category with ratio of 0.37 or network 18 and 20 from visual category with ratio 0.41. We also see negative correlation between different groups like network 20 (visual network) and network 26 (cognitive control) with ratio −0.2.

Furthermore, we also illustrate changes in this FNC over time (i.e., dynamic FNC [43]) to provide additional insight into the model behavior regarding encoding temporal variation. To do this, we computed the cross-correlation between all networks using a sliding window technique with window size of 50 and overlapping ratio of 10. For this analysis, we computed the correlation between all networks within the first 50 timepoints, then we moved the sliding window 40 timepoints forward to repeat the calculation using the next batch of timepoints by shifting the window by 10 timepoints from the previous one. Next, we proceeded with remaining timepoints which resulted in 12 distinct heatmaps, vary over the time and clearly shows dynamicity of functional connectivity network as is shown in figure 10. Given that we now have access to the full 5D space-time-network information, this can be further exploited in future studies to evaluate dynamics over time or space or both.

**Fig. 10.**
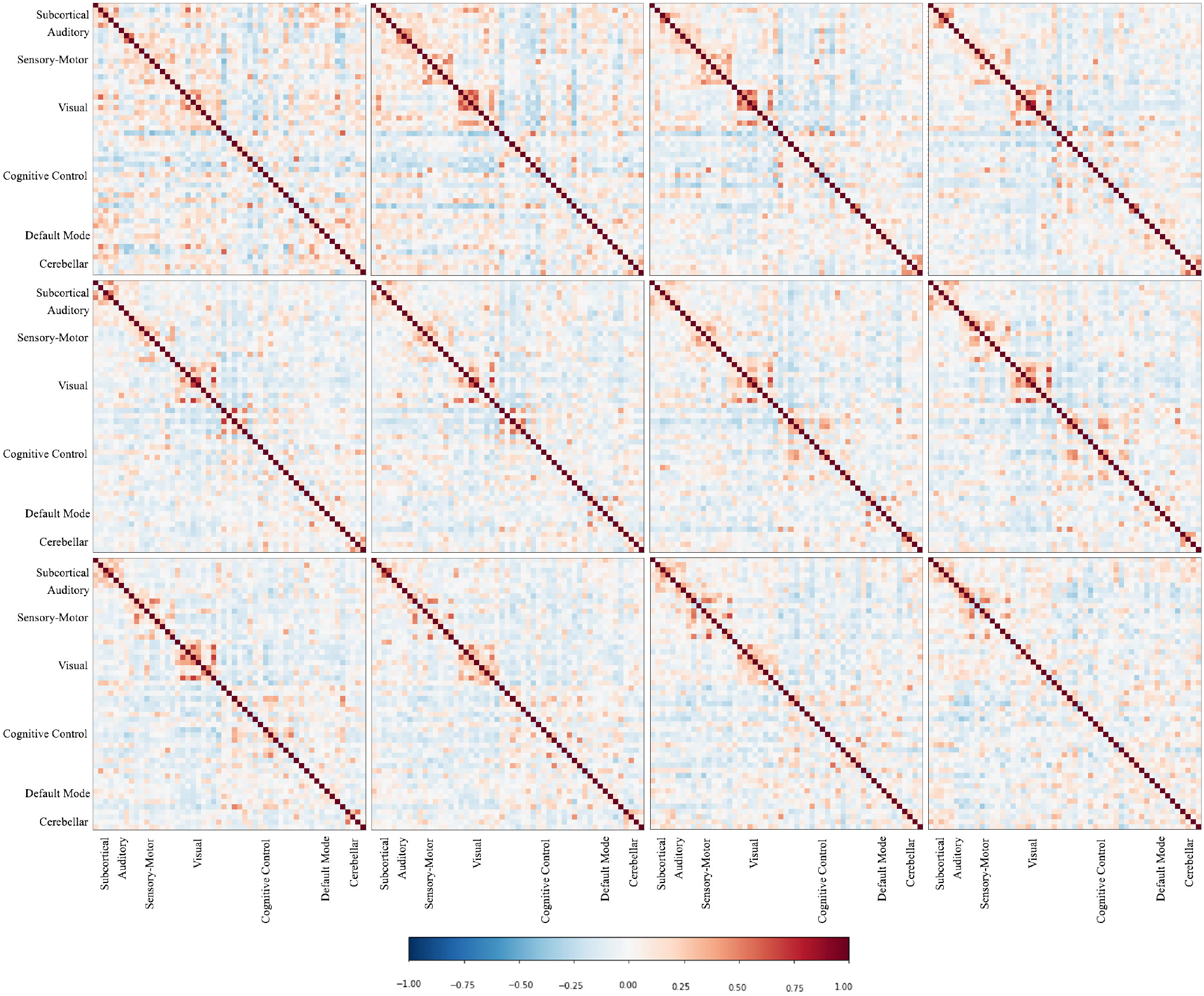
Dynamic FNC is shown by 12 heatmaps computed by sliding window technique for a sample subject and each of heatmaps shows the correlation between all networks for 50 timepoints that overlap 10 timepoints with the previous one ordered left to right and top to bottom. A dynamic FNC approach emphasizes variation of functional connectivity over time which provide an important tool for capturing temporal variation in functional connectivity (i.e., the chronnectome [19]), encoded by our model.

## VI. Discussion

We introduce a novel approach to perform a full 5D parcellation of functional activity, thus preserving the, typically ignored, spatiotemporal behavior of each intrinsic network. We implemented our approach by training a residual network to predict the probability of each voxel belonging to each brain network using ICA priors. Results show our model not only generates reasonable maps of brain segments but is also able to captures spatiotemporal dynamics of the brain, providing a rich feature space useful for further analysis.

Preliminary results show that the proposed model captures differences among subjects, spatial variability within networks, and temporal coupling among different networks. Hence, we see obvious differences in model output for different subjects as is shown in figure 4 for a sample sensory-motor network. Results show that the magnitude of active region vary from subject to subject along with smooth changes in boundaries and shape of active region. Moreover, our approach is able to encode the full spatiotemporal patterns while preserving within and between subject information. This provides a new window into brain activity, highlighting information that is missed by approaches that ignore spatial dynamics. For example, if we focus on the sensory-motor sagittal view of absolute differences plot, we see 3 hot spots located along the SM border that exhibit high variability (temporal fluctuations) over time. This information is not visible in the mean activity, e.g., the original model output. We also see a source of high variability in the auditory network coronal view in the absolute differences plot which is shown in the mean map. This may suggest the network is transiently linking to other network such as the default mode network and should be explored in further studies with larger numbers of subjects. Additionally, exploration of the temporal variation shows high correlation within the same network categories. It is worth mentioning that we observe notable correlation between two different categories (domains) as well, for example, network 7 and 8 from the auditory and sensory-motor domains show high inter-domain correlation, but we also see negative/inverse correlation for some other networks which demonstrate the fact that an increase in activity of a network is linked to a decrease in other network like network 19 from visual category with network 31 from cognitive control which show an inverse activity. We also studied dynamic FNC to evaluate the ability of the model to capture temporal variation in the coupling between networks. Accordingly, we observe significant changes in correlation or anti-correlation between different networks over time as shown in figure 10 and is a consequence of model capability in encoding temporal variations. Together these results highlight the richness of the estimated 5D output and its potential to be applied to study various aspects of the healthy and disordered brain.

A wide range of methods from other research domains have been adapted to study brain connectivity, including the Spatio-Temporal Graph Convolution (ST-CGN) [38] which took advantage of a graph convolutional network to model the non-stationary properties of functional connectivity by learning the importance of edges between brain networks and showed 80 percent accuracy in predicting gender and more than 75 percent on predicting age using the encoded temporal dynamics. However, this model, like most others, did not include spatial variability as a consequence of using a predefined atlas for extracting ROIs. Ignoring spatial and temporal variability may lead to incomplete or even incorrect conclusions and is a motivation for our approach that generates probabilistic maps which not only encode temporal dynamics but also capture spatial variability (e.g., size, shape, translation) between subjects. Similarly, a study of rs-network dynamics employed for schizophrenia diagnosis [44] trained a support vector machine (SVM) with functional connectivity data to classify subjects into patient/control showed 90 percent of accuracy in classification task, however this approach also did not model the spatial or spatio-temporal dynamics. Functional connectivity data has also been used to study Alzheimer’s disease [45] by training an end-to-end deep recurrent neural network with rs-fMRI data. This study resulted in 90 percent accuracy in classification and highlights the importance of spatio-temporal dynamics in biomarker studies, however the end-to-end model also comes with a limitation in that does not naturally enable separation of the mixture of spatial and temporal variabilities. We attempted to address these limitations with our proposed parcellation model which blends together the strengths of deep learning and blind source separation to enhance interpretability with a focus on brain networks. Our proposed model also has a strength relative to typical methods in brain parcellation since it encodes spatio-temporal dynamics into 5D probabilistic maps which can be used for further studies on brain functionality.

## VII. Conclusion

We proposed a novel 5D brain parcellation method using a deep residual network structure which is able to capture spatio-temporal dynamics of the brain. Our method goes beyond existing brain parcellation techniques by providing information about the full spatio-temporal variation over space, time, and networks. We utilized ICA priors to train a model with deep residual structure in order to predict probability of each voxel in a given timepoint belonging to each brain network and obtained a full 5D result after additional post processing steps. According to the results, the proposed approach provides output which is sensitive to individual variations as demonstrated by comparing same network and timepoint for different subjects; Moreover, we studied model performance in differentiating two groups of young and old subjects which showed significant differences in model output between the two groups. We also studied spatial variation via computing timepoint wise differences of model output that reveals regions which simultaneously fall within network boundaries but are different from those of the overall mean activity. This is a key advantage of our approach to provide additional insight into brain activity. Furthermore, we evaluated temporal interactions among networks by computing functional network connectivity (FNC) by grouping all networks into the various domains (subcortical, auditory, sensory-motor, visual, cognitive control, default mode and cerebellar categories) and observed the expected modular output, including generally higher correlation within each domain. We believe that the key feature of the proposed approach in encoding spatio-temporal dynamic of brain networks provide an infrastructure to study brain disorders, cognitive ability and also similarity which are our high-priority future objectives.

